# Rapid and consistent evolution of colistin resistance in *Pseudomonas aeruginosa* during morbidostat culture

**DOI:** 10.1101/080960

**Authors:** Bianca Regenbogen, Matthias Willmann, Matthias Steglich, Boyke Bunk, Ulrich Nübel, Silke Peter, Richard A. Neher

**Author notes:** present address: Institute of Plant Breeding, University of Hohenheim, Stuttgart, Germany (Bianca Regenbogen). joint first authors (Bianca Regenbogen, Matthias Willmann).

## Abstract

Colistin is a last resort antibiotic commonly used against multidrug-resistant strains of *Pseudomonas aeruginosa*. To investigate the potential for *in-situ* evolution of resistance against colistin and map the molecular targets of colistin resistance, we exposed two *P. aeruginosa* isolates to colistin using a continuous culture device known as morbidostat. Colistin resistance emerged within two weeks along with highly stereotypic yet strain specific mutation patterns. The majority of mutations hit the *prmAB* two component signaling system and genes involved in lipopolysaccharide synthesis, including *lpxC*, *pmrE*, and *migA*. In seven out of 18 cultures, we observed mutations in *mutS* along with a mutator phenotype that seemed to facilitate resistance evolution.

---

The surge of multidrug resistance has evolved into a serious complication of modern medicine (Tzouvelekis *et al.*, 2012). Particularly dangerous is *Pseudomonas aeruginosa*, a Gramnegative pathogen known to cause severe infections with high mortality in immunocompromised individuals (Gould and Wise, 1985). Antibiotic drug resistance exacerbates this situation (Kang *et al.*, 2003; Tumbarello *et al.*, 2011). Extensively drug resistant hospital strains are often only susceptible to colistin, which has become an indispensable drug of last resort (Katz *et al.*, 2016).

Colistin belongs to the polymyxin family and has a broad activity against most clinically relevant Gram-negative bacteria. Polymyxin B and polymyxin E (colistin) are currently used for clinical application. Resistance to colistin has been found to be caused by the phosphoethanolamine transferase enzyme MCR-1 located on mobile genetic elements (Liu *et al.*, 2016; Malhotra-Kumar *et al.*, 2016) but has also been reported due to chromosomal mutations (Lee *et al.*, 2016) and could emerge under antibiotic exposure in clinical settings (Jochumsen *et al.*, 2016; Noteboom *et al.*, 2015; Snitkin *et al.*, 2012). Given the potentially severe consequences for patients, it is important to elucidate the underlying evolutionary mechanisms involved in the evolution of colistin resistance.

Recent advances in sequencing technology have made it possible to follow the evolution of bacterial populations over long times and in great detail (Barrick *et al.*, 2009). Evolution experiments are particularly valuable to explore and recapitulate the pathways along which resistance evolves, the nature and order of mutations that arise, and the speed at which resistance emerges. Toprak *et al.* (2012) have presented a detailed study of resistance evolution in *E. coli* under sustained selection pressure for resistance using a custom made device called *morbidostat*. The morbidostat continuously adjusts the concentration of antibiotics to maintain a constant growth rate of bacteria in stirred liquid culture. The bacteria are challenged just enough that they still grow, but are under strong pressure to evolve resistance. By sequencing the evolving *E. coli* populations, Toprak *et al.* (2012) showed how mutations accumulated and affected antibiotic resistance.

Here, we used a customized morbidostat setup to study the evolution of colistin resistance in two clinical *P. aeruginosa* isolates. All cultures from both strains evolved resistance within two weeks and increased their tolerance to colistin in liquid culture about 100 fold. On plates, the increase in the minimal inhibitory concentration (MIC) was less pronounced. In agreement with previous characterizations of colistin or polymyxin resistance in *P. aeruginosa* and other bacteria (Moskowitz *et al.*, 2012, 2004; Olaitan *et al.*, 2014), we observed the rapid emergence and spread of diverse mutations in *pmrB* and other genes involved in lipid A and lipopolysaccaride synthesis. The evolution of resistance sometimes went along with mutator phenotypes (due to mutations in *mutS*), which increased mutation rate by approximately 100 fold.

## Results

### Characteristics of patient isolates

Two clinical isolates (PA77 and PA83) were investigated, which originated from two different patients with *P. aeruginosa* bloodstream infections. Both strains exhibited extensively drug resistant phenotypes (Magiorakos *et al.*, 2012), being non-susceptible to all antibiotics except colistin (PA77 and PA83) and fosfomycin (PA77). Multilocus sequence typing revealed PA77 belonged to sequence type ST308 and PA83 to ST233 (Willmann *et al.*, 2014).

### Whole genome sequences of patient isolates

We sequenced the strains PA77 and PA83 with 98x coverage using PacBio long read sequencing technology. Together with high-fidelity short reads from the Illumina HiSeq platform, we were able to assemble one circular chromosome of length 6.82Mb and one 398kb plasmid for strain PA83, while the assembly of strain PA77 resulted in three contigs of size 3.7Mb, 2.3Mb, and 1Mb and one circular plasmid of size 40kb. The plasmid of PA77 had consistently 2-3 fold higher coverage than the chromosome, suggesting it is present in multiple copies.

The 40kb plasmid of PA77 contained several resistance genes (incl. *blaIMP-8* as identified by resFinder, (Zankari *et al.*, 2012). The majority of resistance genes (incl. *blaVIM-2*) of PA83 reside in the chromosome. ResFinder results for both strains are available as supplementary Tables S1 and S2.

### In-vitro resistance evolution against colistin

We performed three replicated experiments (two for strain PA77 with four and five parallel cultures, respectively, and one experiment for strain PA83 with nine parallel cultures) selecting for colistin resistance mutations in a modified morbidostat set-up (Toprak *et al.*, 2013). Our morbidostat can culture 15 populations in parallel. We used a culture volume of 20ml LB media and maintained cultures at an optical density around 0.1 OD600, corresponding to about ~ 4 × 10^8^ bacteria per culture (Kim *et al.*, 2012). The optical density of each culture was monitored every 30 seconds. Every ten minutes, a computer program calculated the rate at which the bacteria grew and colistin concentration in the vials was increased or decreased by adding concentrated colistin solution or medium, respectively. The decisions to increase or decrease the concentration were made automatically by the computer program as described in Materials and Methods.

In contrast to the antibiotics used by Toprak *et al.* (2012), colistin is a bactericidal antibiotic resulting in less stable feedback on growth. Sometimes, we observed sudden population collapse when the colistin concentration increased by small amounts. These sudden collapses were only observed during the first ten days before substantial resistance emerged.

We will focus here on the second experiment with strain PA77, where five parallel cultures (vials v01 - v05) were selected for colistin resistance. The first experiment with strain PA77 (described in the supplement) delivered similar results but ran for only 15 days. The experiment with strain PA83 (9-fold replicated) also showed similar patterns of evolution. However, frequent mutator phenotypes (see below) and erroneous concentrations of colistin stock solutions used for six of the 22 days of this experiment make this run less interpretable. In each of the experiments, we took samples three times a week for deep sequencing, plated cultures for purity control, and performed Etests to assess resistance against colistin on plates. In parallel, we inferred the concentration of colistin in the liquid cultures from the known schedule of colistin additions and dilutions for each vial. Both resistance measurements are shown in Figs. 1 and 2.

**FIG. 1.**
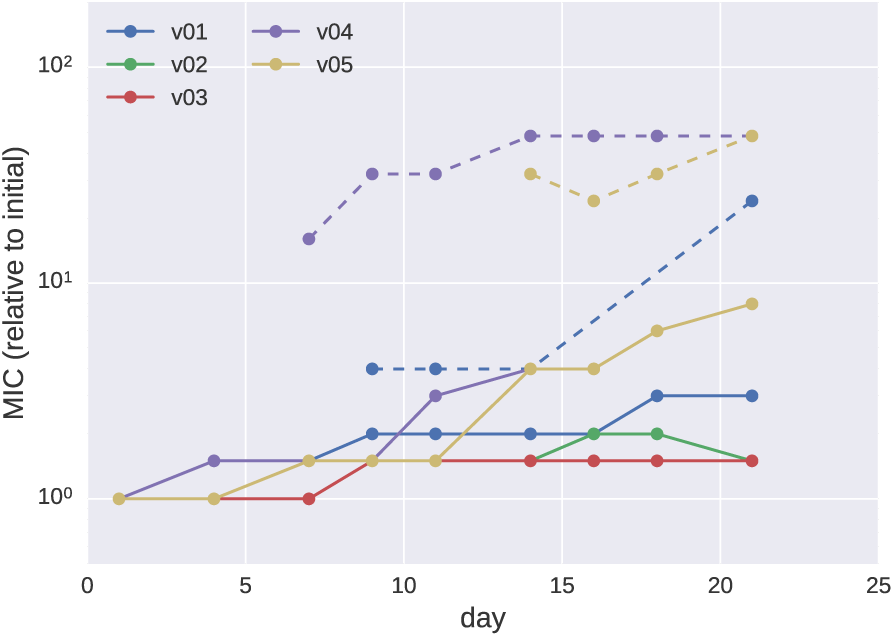
Etest results for PA77. MICs determined in Etests increased moderately over the course of the experiments (v01 v05 denote the five culture vials for strain PA77). Sub-populations showing a higher MIC than the main population were observed in some vials after 7 days and are shown as dashed lines.

**FIG. 2.**
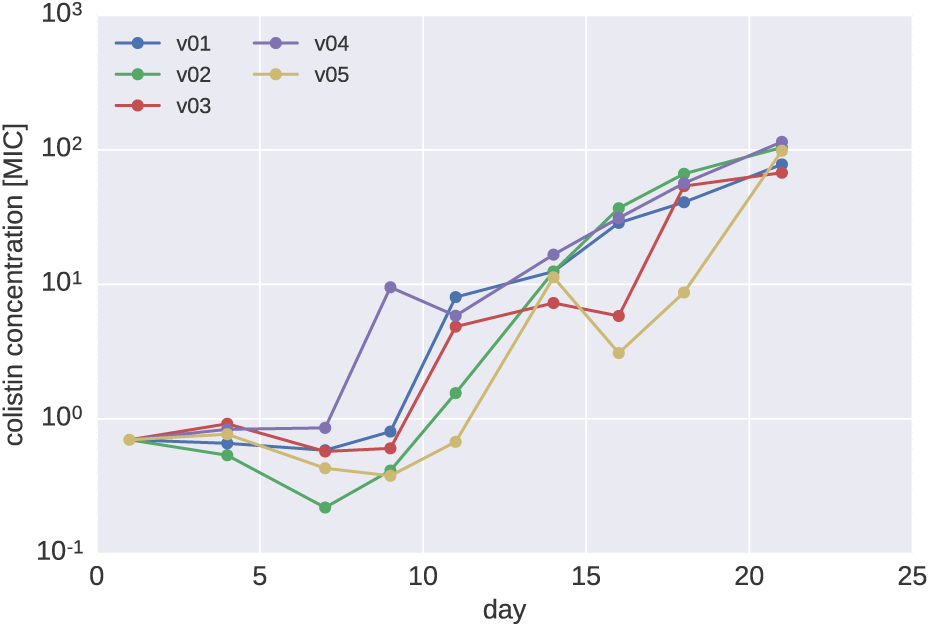
Colistin concentrations during morbidostat culture of PA77. The colistin concentration necessary to inhibit growth in the morbidostat increased much more dramatically than MICs measured in Etests. Colistin concentrations are given in units of the MIC of the initial cultures (2.8*µg/ml*).

The colistin concentration that was tolerated in liquid culture increased by about 10-fold between day 7 and 12 and further increased by 10-fold towards the end of the experiment (Fig. 2). Similar, but less pronounced increased colistin tolerance was observed in Etest measurements on plates (Fig. 1). In addition to 2-10 fold increase of the colistin MIC of the bulk population, the evolved populations contained sub populations that tolerated even higher colistin concentrations as measured by Etests (dashed lines in Fig. 1). These subpopulations, which grew as morphologically smaller colonies, arose for the first time after 7 days of colistin treatment with strain PA77.

**TABLE I.** Mutations repeatedly observed in cultures v01 to v05 of strain PA77. The full list including annotation of each mutation is available as supplementary Table S3.

| Gene locus tag | v01 | v02 | v03 | v04 | v05 |
| --- | --- | --- | --- | --- | --- |
| *pmrB* PSA77_03611 | V9A,L17Q | E320K,L90Q | V9A | P216Q | P254L |
| *pmrE* PSA77_06074 | Y28N | Y28C | Y28N | Y28C | Y28N |

### Convergent evolution in genes involved in LPS synthesis

Mutations in genes that mutated in at least two cultures and became a majority variant are summarized in Tab. I. All cultures developed mutations in *pmrB*, one of which (V9A) was observed both in culture v01 and v03. These mutations presumably result in constitutive activity of *pmrB* (Moskowitz *et al.*, 2012). Besides mutations in *pmrB*, strain PA77 developed repeated mutations in an UDP-glucose-6-dehydrogenase (*pmrE*, also known as *udg*). In this case, the same codon was hit multiple times, resulting in two Y28C and three Y28N mutations. The gene *pmrE* is known to be a tyrosine phosphorylation target and has been implicated in colistin resistance evolution (Lacour *et al.*, 2008). Interestingly, position 28 is a cysteine in the majority of *P. aeruginosa* reference genomes including PA83, which might explain the absence of *pmrE* mutations in PA83.

In addition to the recurrent mutations in *pmrB* and *pmrE*, two cultures of our preliminary experiment with PA77 evolved mutations in *imp/ostA*. *Imp/ostA* has been described as an essential outer membrane protein (Braun and Silhavy, 2002). Mutations in *imp/ostA* result in increased membrane permability and organic solvent tolerance (Aono *et al.*, 1994; Sampson *et al.*, 1989). A full list of all mutations observed in PA77 at frequencies above 25% is provided as Table S3.

Strain PA83 repeatedly mutated UDP-3-O-[3-hydroxymyristoyl] N-acetylglucosamine deacetylase (*lpxC*), which participates in the biosynthesis of lipid A and could, therefore, be involved in the progression to colistin resistance as shown in *P. aeruginosa* (Jochumsen *et al.*, 2016) and *Acinetobacter baumannii* (Moffatt *et al.*, 2010). The protein was mutated in every culture and suffered from multiple mutations in 5 out of 10 vials. All mutations were nonsynonymous and the V223A and A108T mutation were shared among 3 cultures. In addition to *lpxC*, all cultures developed mutations in *pmrB*, three out of nine mutated *pmrA*, and seven out of nine mutated alpha-1,6-rhamnosyltransferase *migA*, which is thought to be involved in the outer oligosaccharide synthesis (Poon *et al.*, 2008). Additional recurrent mutations were observed in *lpxO2*, an asparagine synthase, a putative acetyl transferase, and several other genes. A full list of all mutations observed is provided as Table S4. Overall, many more mutations arose in PA83 than in PA77. This excess of mutations is partly explained by the frequent rise of mutator phenotypes (see below).

### Dynamics of mutations

Deep population sequencing (mean coverage *>*150x) of the continuously cultured populations in the morbidostat allowed us to study the dynamics of mutations in the entire genome and to quantify the competition between different lineages.

During experiments with PA77, the first major increase in colistin tolerance was observed between day 7 and 10, concomitant with a quick rise of mutations in *pmrE* and *pmrB* (see Fig. 3). *pmrB* mutations tend to occur first, followed by mutations in *pmrE*. In two of the PA77 cultures (v01 and v02), two *pmrB* mutations arose, and the initially successful mutant was later replaced by the other one that also carried the mutation in *pmrE*. In culture v05, a transient mutation in *prmA* was observed, which was outcompeted by a lineage carrying mutations in *pmrB* and *pmrE*. In cultures v01, v02, v03, and v04 additional mutations rose to intermediate frequencies during the last few days of the experiments (see Fig. S1), possibly explaining the increase in colistin tolerance during the second half of the experiment. The frequency trajectories of all sweeping mutations are provided in supplementary Tables S5, S6, and S7.

**FIG. 3.**
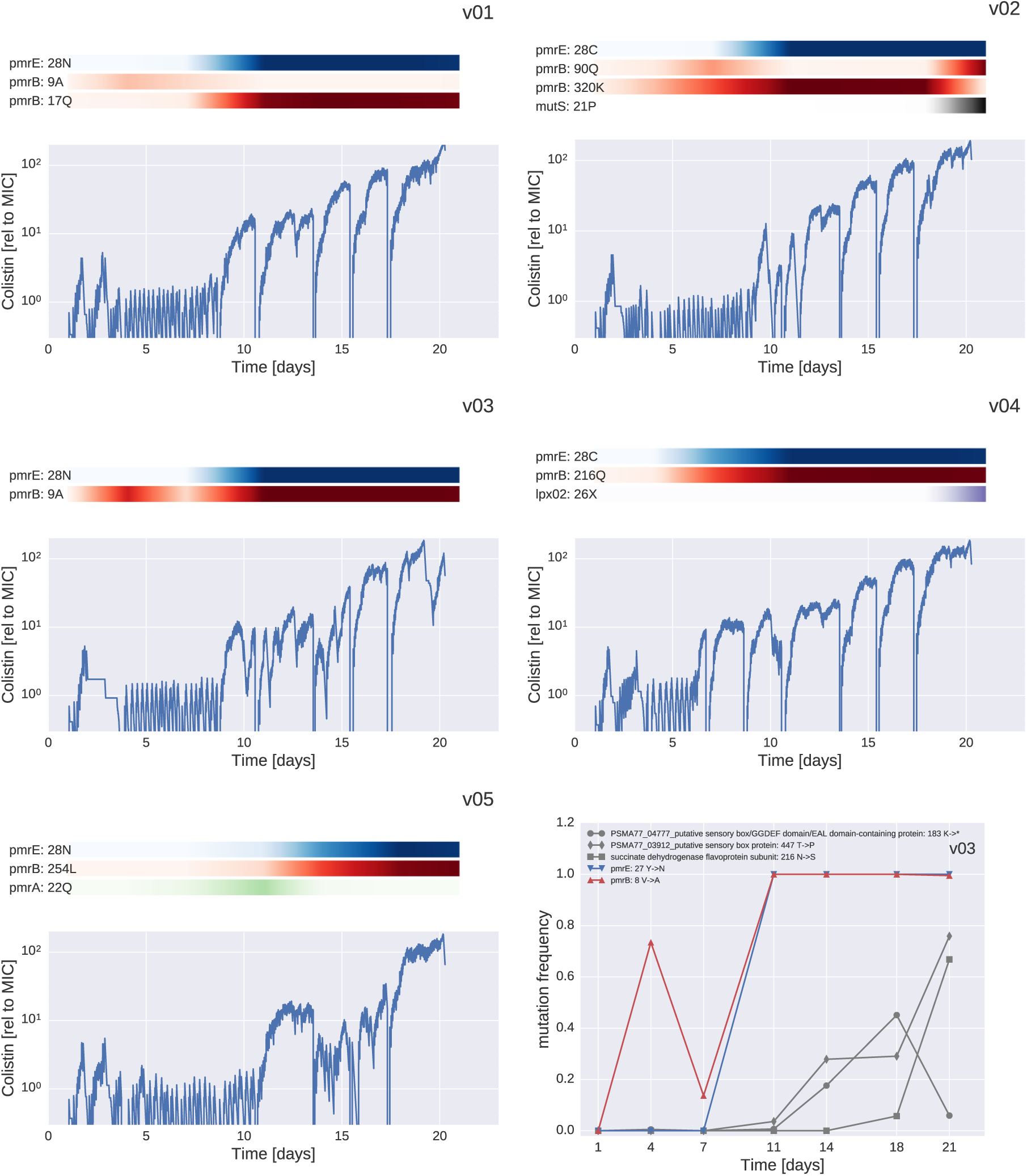
Resistance evolution in PA77. For each culture vial, the plot shows the dynamics of colistin concentration in liquid culture. This concentration is inferred from the cycles of colistin addition and waste removal in 10 minute intervals. The shaded bars above the plots show the abundance of different mutations during the experiment. The frequencies of *pmrE* and *pmrB* mutations correlate well with colistin tolerance. The deep dips in colistin concentration every 2-3 days correspond to transfers to fresh culture vials and mark the time points at which samples were taken. Additional mutations in other genes, mostly specific to individual cultures rose to intermediate frequencies towards the end of the experiment, as shown in the bottom right panel for culture v03 (for analogous plots for other cultures, see Fig. S1).

In experiments with PA83 more complicated dynamics of mutations in three commonly mutated genes (*pmrA*, *pmrB*, and *lpxC*) were observed. In many cases the same gene was mutated independently several times at different positions and apparently functionally similar sub-populations compete. This higher diversity is most likely related to the almost ubiquitous mutator phenotypes observed (see below and supplementary materials). Trajectories of mutations observed in PA83 are shown in Fig. S7 and S8.

#### Preexisting variation

A number of loci were already polymorphic in the initial samples and the frequencies of these preexisting mutations changed over time as new adaptive mutations arose and the population composition changed (Fig. S1, S4, S7, and S8). In the majority of cultures a particular subpopulation came to dominate. In PA77, the more successful subpopulation carried a P282S mutation at locus PSA77 01281, annotated as *putative pseudouridylate synthase*. The variant was initially at 40% frequency and was at frequencies *>* 90% in the final sample in 8 out of 9 cultures. In PA83, all populations fixed the full length allele of the sensory histidine kinase *CreC* (locus PSMA83 00508), even though 75% of the initial population had a premature stop at codon 319.

### Mutator phenotypes

The culture v02 of PA77 developed a mutator phenotype and carried the mutation H21P in *mutS*. This mutation rose rapidly in frequency between days 17 and 22 at the end of the experiment. In the last sample, 42 mutations were observed at high frequencies that were not apparent earlier. Even though we lack information on the linkage between these mutations, the most likely explanation is that they arose quickly after the mutation in *mutS* and were carried to high frequency through linkage with a mutation that conferred a benefit in the culture system. The *mutS* mutation might have been around for many days before it became frequent and other mutations will have accumulated throughout this time. The full list of all observed mutations can be found in Table S3, but mutations that likely arose after mutations in *mutS* are omitted from the graphs.

Mutator phenotypes were much more common in PA83, where dozens of mutations were observed in seven out of nine cultures all of which carried the mutation T51P in *mutS*. This mutation might have preexisted at low frequency in the initial population and rose in frequency repeatedly because it facilitated resistance evolution. The majority of mutations that rose along with T51P in *mutS* were unique to each culture, suggesting that these mutations accumulated after the different cultures were inoculated.

In cultures with *mutS* mutations we observed between 30 and 100 mutations (Tables S3 and S4) above 20%, the majority of which were observed only in one culture and thus likely arose during the 21 days of culture in the morbidostat. Fig. 4 shows the number of synonymous and intergenic mutations vs the number of non-synonymous mutations observed in the last sample. Each mutation is weighed by its frequency in the population. In contrast to mutations in non-mutators, which are almost exclusively within coding regions and result in amino acid differences, about 1/3 of mutations in mutator strains are synonymous or intergenic. This is consistent with most of these additional mutations being a random byproduct of elevated mutation rates in the mutator strains.

**FIG. 4.**
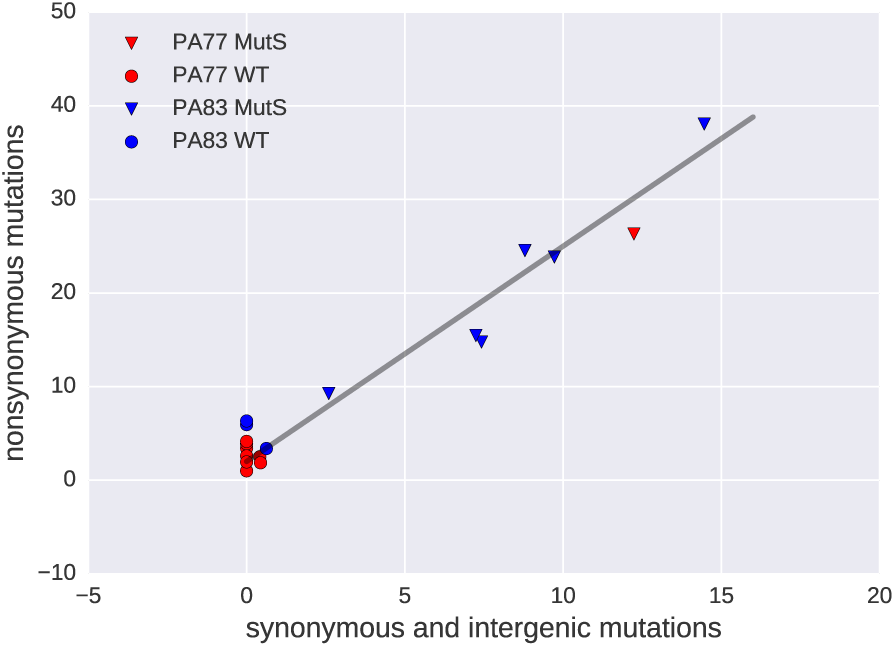
Mutation patterns in mutators and non-mutators. In cultures with no mutations in *mutS*, only nonsynonynous mutations are observed. Most populations carrying a mutation in *mutS* accumulated the expected mix of synonymous and non-synonymous mutations. Mutations are weighted by their frequency in the final sample.

In total, we observed, 43 and 152 frequency weighted synonymous and non-synonymous mutations after a total of 154 days of culture in the seven cultures with a *mutS* mutation. The overall mutation rate of mutator strains was, therefore, about 1.3 one mutations per day. The morbidostat dilution forces a doubling time of 90 min. Hence the observed mutation rate corresponds to about 0.08 mutations per replication. Given a genome size of 7Mb, the mutation rate of mutator strains is on the order of 10^*−*^8 per site and generation.

Only synonymous mutations are suitable to compare the mutation rates of mutator and non-mutator, since most nonsynonymous mutations in non-mutators are likely adaptive in presence of colistin. In cultures without mutations in *mutS*, the frequency weighted number of synonymous mutations is 1.1 in 242 days of culture, while cultures with mutators accumulated 43 synonymous mutations in 154 days. These data suggest that the mutation rate of the *mutS* mutants is increased by approximately two orders of magnitude relative to wildtype, consistent with previous results (Chopra *et al.*, 2003; Taddei *et al.*, 1997).

### Copy number variations

A small number of deletions were observed during resistance evolution. Two prominent almost adjacent deletions occurred in culture v05 of PA77 (Fig. 5) and partially deleted the metallo-beta-lactamase (MBL) IMP-8 and an aminoglycoside 3’-phosphotransferase (aph(3’)-XV). The regulation of other resistance proteins in the vicinity might also have been affected. The loss occurred between day 11 and day 18 in parallel with the spread of mutations in *pmrE* and prmB. The disappearance of the IMP-MBL has been confirmed by PCR and the breakpoints in coverage are both supported by split reads. The deletions occurred at identical positions at the end of a duplicated stretch (18666-19310 and 20647-21291 on the plasmid containing *aacA4*).

**FIG. 5.**
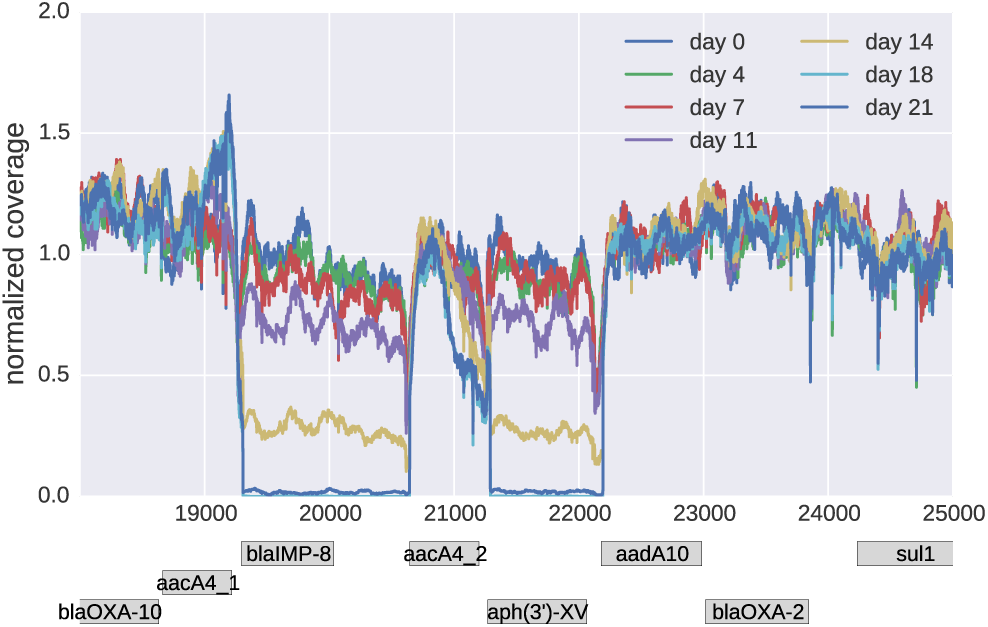
Loss of resistance genes. One of the cultures (v05, PA77) lost two neighboring chunks on the plasmid gradually between day 11 and day 18. The block between 20647-21291 corresponds to a duplicated sequence also found at positions 18666-19310. Both breakpoints are confirmed by about 50% of reads suggesting a bona-fide double deletion.

While PCR was still positive for IMP at day 14, it turned negative at day 21. Of note, resistance to meropenem and aminoglycosides remained unaltered in PA77 at day 21, suggesting alternative mechanisms that were responsible for the observed phenotype. Preservation of aminoglycoside resistance could have been mediated by additional resistance conferring genes that are part of the PA77 plasmid (*aacA4*, *aac(6’)Ib-cr*, and *aadA10*, see Table S1) and that were not deleted. The preservation of meropenem resistance is less conclusive. All OXA-enzymes found in PA77 do not hydrolyze carbapenems (Evans and Amyes, 2014). However, a number of efflux systems that have been described to cause meropenem resistance when overexpressed have been identified in PA77 (see supplement) and could explain the unaltered resistance phenotype (Rodr´ıguez-Mart´ınez *et al.*, 2009).

### Discussion

The morbidostat continuously adjusts drug concentration such that bacteria are always challenged to evolve higher tolerance against the drug while still being able to grow (Toprak *et al.*, 2012, 2013). In contrast to transposon knock-out screens for polymyxin resistance (Fernández *et al.*, 2013), direct selection for resistance by the morbidostat and whole genome deep sequencing allows the unbiased detection of mutations associated with resistance (Punina *et al.*, 2015). Furthermore, compared to classical experimental setups to investigate antibiotic resistance evolution such as serial dilution protocols or chemostats, the morbidostat approach allows a higher degree of replication (Jansen *et al.*, 2013; Jochumsen *et al.*, 2016). Multifold replication is essential to quantify convergent evolution, prevalence of different evolutionary pathways, and the mutational target of resistance.

The continuously increasing but sub-lethal antibiotic concentrations in the morbidostat might simulate a clinical situation in which the antimicrobial agent does not reach lethal or inhibitory quantities in all compartments of the infection. Resistant bacteria might evolve in such insufficiently suppressed compartments and gradually spread while more resistance mutations accumulate. *Baym et al.* (2016a) have recently demonstrated that the kinetics of drug resistance evolution along spatial gradients is similar to the kinetics observed in morbidostat. While the morbidostat is not intended as a faithful model of the situation *in-vivo*, it is nevertheless useful to determine a bacterial populations’ capacity to spontaneously evolve resistance or to select already existing rare resistant mutants in a clinical setting. We have focused on colistin resistance since the recent increase in colistin use against XDR pathogens greatly increased the potential for resistance evolution against this last resort drug. Understanding the pathways and kinetics of colistin resistance evolution is of paramount importance.

Colistin tolerance in liquid culture increased 10-fold within approximately 10 days and 100-fold after 20-days in a bacterial population of ~4 ×10^8^. MICs, as measured by Etest on plates, also increased consistently but only by about 2-fold and 4-fold after 10 and 20 days, respectively. However, much more resistant sub-populations were visible on Etest plates in a fraction of the cultures after about 10 days. The discrepancy between liquid culture and Etest measurements of colistin tolerance could either be explained by the nature of the tests or a shift in the population composition that might have occurred while preparing cultures for Etests. Agar dilution, disk diffusion, and gradient diffusion have been reported to be problematic, and EUCAST has released a warning suggesting to only use broth microdilution for diagnostic testing at the moment (EUCAST, 2016).

Selection for colistin tolerance resulted in a reproducible rise of mutation in *pmrB* and *pmrE* in strain PA77 and *lpxC* and *pmrAB* in PA83. The *pmrE* mutations Y28C reverted the position to the amino acid shared by the majority of *P. aeruginosa* reference genomes in NCBI. Since the strain PA77 is a clinical isolate with a complicated history of antibiotic exposure, it is not clear whether this mutation is a reversion of a previously adaptive mutation or a mutation that is specific to colistin resistance in the genetic background of PA77. Tyrosine phosphorylation of *pmrE* has been implicated in colistin resistance (Lacour *et al.*, 2008). *pmrA* and *pmrB* mutations have been previously reported to mediate colistin resistance (Moskowitz *et al.*, 2012, 2004; Olaitan *et al.*, 2014; Pamp *et al.*, 2008). Most other mutations that arose repeatedly are involved in lipopolysaccaride synthesis and lipidA biosynthesis – as expected in case of colistin. Jochumsen *et al.* (2016) recently reported mutations that arose during colistin resistance evolution during serial transfer to the laboratory strain PAO1. They also found parallel mutations in genes *pmrB* and *lpxC*. However, other loci that frequently mutated in experiments by *Jochumsen et al.* (2016), such as PA5194 (8 out of 9) and PA5005 (5 out of 9), rarely mutated in our experiments. We found two mutations in homologs of PA5194 (one in v02 of PA77 at locus PSA77 04096, one in v03 of PA83 at locus PSMA83 05923) and no mutation at in *opr86* or at locus PA5005. We did not observe a strict order in which the mutations arose. While *pmrB* tends to be the first mutation that rises to high frequency, mutations in *pmrA* and *lpxC* preceeded *pmrB* in cultures v05 of PA77 and v02, v06 and v12 of PA83.

The observed *in-vitro* loss of the plasmid-encoded genes blaIMP-8 and aph(3’)-XV in PA77 is particularly interesting since it might not only mirror vivid dynamics in genomic rearrangements under antibiotic exposure but could also constitute an example of the previously proposed collateral sensitivity concept (Baym *et al.*, 2016b; Imamovic and Sommer, 2013). This suggests that bacteria that develop resistance to one antibiotic could become more susceptible to another antibiotic at the same time. Loss of antimicrobial resistance genes against antibiotics to which the strain was not currently exposed, and the simultaneous emergence of colistin resistance would fit this concept. However, we have not observed changes in the aminoglycoside or carbapenem susceptibility pattern of PA77 after the loss of these genes, most likely due to alternative resistance traits. Collateral sensitivity in extensively drug resistant pathogens might be a phenomenon that heavily depends on a strain’s overall resistance potential and might be hard to predict without full characterization of all resistance factors (e.g., by whole genome sequencing) (Buchanan *et al.*, 2014). Comparison of the mutations observed in PA77, PA83, and evolution experiments by (Jochumsen *et al.*, 2016), suggests strain specific mutational pathways to colistin resistance with a common core of *pmrB* and to a lesser extend (**?**). Both genes have a large mutational target and many different mutations seem to contribute towards colistin resistance (Olaitan *et al.*, 2014).

Our results underscore the potential importance of mutator phenotypes in bacterial resistance evolution, as also reported by (Jochumsen *et al.*, 2016). Once the mutation rate is high, drug resistance mutations are much more rapidly discovered in the mutator lineage, in particular when multiple mutations are necessary to convey full resistance (Chopra *et al.*, 2003; Marvig *et al.*, 2013; Taddei *et al.*, 1997). Our estimate of the mutation rate suggests that in cultures dominated by mutator strains, most mutations are produced every generation (the product of mutation rate and population size exceeds 1), while in absence of mutator alleles a specific mutation would take a few days to be discovered. However, the mutational target is much larger than a single site – in particular in *pmrA/B* – such that resistance evolved rapidly even in cultures with no dominant mutator phenotype.

One reason why colistin resistant strains do not emerge as frequently in clinical settings compared to *in-vitro* experiments could be fitness costs and impaired virulence associated with resistance. Lee *et al.* (2016) observed rapid reversion of colistin resistance mutations in *P. aeruginosa*. Similarly, (Fernández-Reyes *et al.*, 2009; López-Rojas *et al.*, 2011) describe fitness costs of colistin resistance in *Acinetobacter baumannii* isolates. Membrane modification associated with colistin resistance might reduce clinical invasiveness, possibly due to a lower attachment ability to host epithelium cells, resulting in a lower colonization potential and a “flushing away” from the invasion site.

The highly parallelized morbidostat approach enables to determine various independent evolutionary trajectories of resistance development. One limitation of our study is that we have investigated only two clinical strains and are, thus, not able to draw general conclusions about a potentially common timeline of mutations towards colistin resistance. Investigation of many clinical strains might reveal common trajectories and high risk mutations that do not yet cause clinical resistance but predispose a strain to become fully resistant. Such pre-resistance marker could be clinically useful to decide whether a combination treatment if possible, especially with aminoglycosides should be started that would have been otherwise avoided due to concerns about cumulative toxicity. This way, findings from *in-vitro* evolutionary experiments could provide knowledge that can be translated into routine diagnostics and treatment, eventually improving our therapeutic concepts and patient care.

## Materials and Methods

### Bacterial strains

The two *P. aeruginosa* strains, ID 77 and ID 83, where isolated at the Department of Haematology in Tèbingen from January 2010 to December 2013 from adult patients with haematological-oncological conditions, such as leukaemia, lymphoma, and multiple myeloma (Willmann *et al.*, 2014). Both isolates were isolated from the patients’ blood during bacteremia.

### Risk assessment

Since clinically relevant pan-resistant strains can potentially emerge during the experiments performed here, risk of the such studies need to be considered carefully. We consulted the local research ethics committee of the University of Tèbingen, which did not express any concerns regarding the study design (reference number: 677/2013BO1) but recommended the consultation of the head of the hospital infection control team to ensure high laboratory safety standards. Subsequently, a standard hygiene protocol for the handling of XDR strains in the morbidostat has been implemented and laboratory personnel has been trained to strictly comply with these practises.

The morbidostat is a closed system and all waste products were immediately deactivated in a container with disinfectant. The *P. aeruginosa* strains used pose little risk to immunocompetent people, and laboratory personnel were not involved in any form of patient care. The experiments were performed in a dedicated room in a research facility.

### Morbidostat and experimental procedures

The morbidostat system was build following the detailed instructions by Toprak *et al.* (2013) with the following modifications: we used (i) DC pumps instead of AC pumps arranged in a different geometry, (ii) an Arduino mega256 microcontroller instead of the MC DAQ card, and (iii) a custom python control software instead of the Matlab based software provided by *Toprak et al.* (2013). The custom written control software is available at github.com/neherlab/python_ morbidostat. In addition, the outlets of the three separate pumps for each culture vial were combined such that only one tube runs from the pump array to each culture vial inside the incubator. The culture volume of each vial was 20 ml and the target optical density was 0.1. Cultures were kept in an incubator at 37^*°*^C.

Before each experiment, the setup was sterilized as suggested by Toprak *et al.* (2013). The initial colistin concentration or minimal inhibitory concentration (MIC) of the strains was inferred by cultivating them at several different colistin concentrations slightly above and below the MIC. The MIC in liquid culture was found to be 2.8 *µg/ml* and 4.8 *µg/ml* for strains PA77 and PA83, respectively.

All morbidostat experiments were started with 7x and 25x MIC concentrations in the colistin reservoirs. As *P. aeruginosa* population developed resistance, the colistin concentrations in the reservoirs was increased such that growth could be regulated by addition of colistin solution from these reservoirs.

The morbidostat recorded the optical density in each vial every 30 seconds. After 10 minutes, the growth rate of each vial was calculated. Depending on the rate of growth and the optical density in the vial, either pure medium was added to dilute the culture and increase growth, or colistin solution (low or high concentration) was added to inhibit growth. The target growth rate was set to a doubling time of 90 min and we used a target OD of 0.1. Fig. 6 shows a flow chart detailing the conditions used to determine whether culture is diluted with medium or colistin solution. Waste products were automatically removed using a 16-channel peristaltic pump and immediately transferred into inactivating solution.

**FIG. 6.**
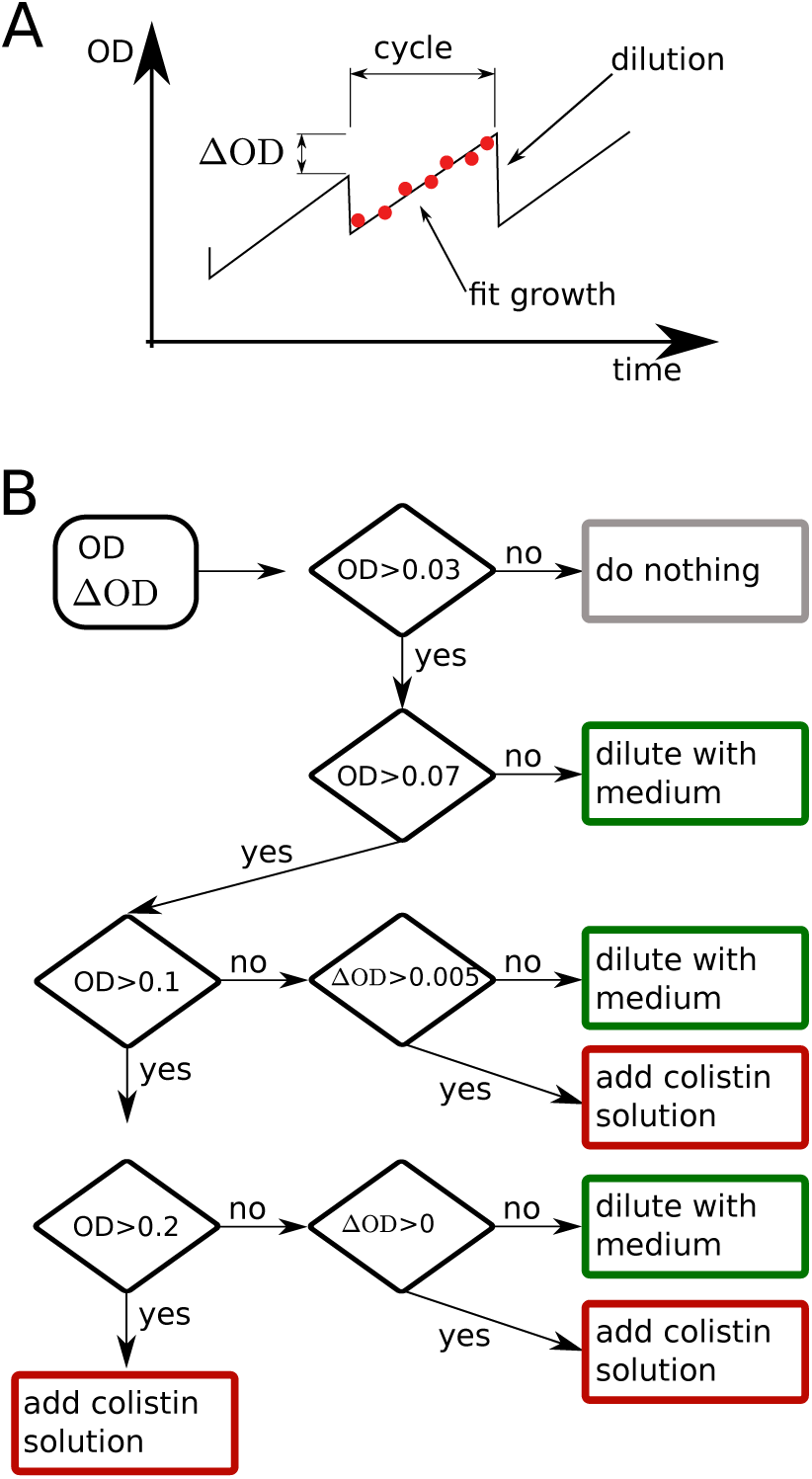
Growth feedback by the morbidostat. A) The morbidostat measures the OD 18 times during a cycle of 10 minutes. At the end of each cycle, the culture is diluted and excess liquid removed. B) The decision whether to dilute the culture and whether to dilute the culture with medium or colistin solution is based on the current OD and the increase of OD (∆OD) compared to the previous cycle.

The morbidostat was programmed to use the colistin solution with the higher concentration whenever the colistin concentration in the respective vial exceed in 1/3 of the colistin solution with the lower concentration. In addition, we limited the rate of colisitin increase to 10% over one hour to prevent too rapid feed back followed by population collapse.

*Sampling.* Every 2-3 days, 1 ml of the bacterial culture was transferred to 19 ml of fresh LB medium in a sterile vial with a magnetic stir bar. To avoid contamination, the vial lid with the inlet was screwed onto a sterile empty vial during the transfer procedure. From every vial 500 *µl* suspension was mixed with 250 *µl* Glycerine 50 % and stored at -80C^*◦*^. To assess purity and to conduct resistance testing with Etests, 10 *µl* suspension was spread on blood agar plates and grown over night at 37 degrees.

*Etest.* Bacterial material was taken from a blood agar plate and diluted with physiological NaCl-Solution to 0.5 McFarland corresponding around 108 CFU/ml. The mixture was plated on Mueller-Hinton agar plates. Colistin Etest strip (bestbion, Cologne, Germany) was placed in the middle of the plate (Andrews, 2001). The bacteria were cultivated over night. After 22 hours, the resulting MIC could be checked. Colonies showing MIC ≥ 4*µg/ml* are classified as clinical resistant limiting treatment options according to EUCAST criteria (Leclercq *et al.*, 2013).

*PCR assays and DNA sequencing.* For the detection of *bla*IMP genes, a PCR amplification was conducted according to a previously described protocol (Pitout *et al.*, 2005). Multilocus sequence typing (MLST) of both strains was performed according to the instructions on the *P. aeruginosa* MLST website (pubmlst.org/paeruginosa/). Sequencing of internal fragments of seven housekeeping genes (*acsA*, *aroE*, *guaA*, *mutL*, *nuoD*, *ppsA* and *trpE*) was done to determine the sequence type.

## Whole genome sequencing and analysis

### Reference genomes

Reference genomes of the PA77 and PA83 strains were determined by PacBio long read sequencing. *P. aeruginosa* DNA was isolated with the *MoBio Ultra Clean Microbial DNA Isolation Kit* according to the manufacturers instructions including the optional RNAse step. DNA from each strain was sequenced in two SMRT cells on a PacBio RSII instrument. For the two genomes, 77,931 and 99,623 PacBio reads with mean read lengths of 15,479 and 11,143 basepairs were assembled using the HGAP.3 protocol implemented in SMRT Portal version 2.3.0. Illumina reads *>*180-fold coverage were mapped onto the assembled sequence contigs using BWA (Li and Durbin, 2009) to improve sequence quality. Annotation was performed using Prokka 1.8 software (Seemann, 2014) and manually supplemented. Genome sequences were submitted to GenBank (http://www.ncbi.nlm.nih.gov/genbank) and assigned accession numbers CP017293 (PA83, chromosome), CP017294 (PA83, plasmid), MJMC00000000 (PA77).

### Population sequencing

For DNA extraction, 10 *µl* thawed sample suspension was plated on blood agar and grown overnight at 37 degrees. Bacterial DNA was isolated by using the *MoBio Ultra Clean Microbial DNA Isolation Kit* according to the manufacturers instructions.

Sequencing libraries of the PA77 samples of the first experiment were prepared by using a modified *Nextera XT* protocol (for details see Zanini *et al.* (2016)) and sequenced to an average coverage of 33x on MiSeq (three chromosomal contigs: 24x, 27x, 24x, plasmid: 56x) with a v2 2x250 bp paired-end kit.

Sequencing of samples from the two later experiments (PA77 and PA83) were prepared with the *TruSeq nano* kit by Illumina and sequenced on a HiSeq 2500 in 2x100bp paired end run. In total we sequenced 35 samples of strain PA77 and 61 samples (three chromosomal contigs: 182x, 212x, 191x mean coverage, plasmid: 457x) of PA83 (chromosome: 175x, plasmid: 388x) on four lanes. Sequencing reads have been submitted to the European Short Read archive and will be available under study accession PRJEB15033 (sample accessions for samples from PA77a, PA77 and PA83 are ERS1284627-ERS1284650, ERS1284651-ERS1284684 and ERS1284685-ERS1284744, respectively).

### Bioinformatic pipeline

*Trimming. TrimGalore!* was used for adaptor clipping and quality trimming with a Phred score cut-off at 20 of the pairedend reads (Krueger, 2015). Resulting fastq-files were checked using *FastQC* (Andrews, 2015).

*Mapping.* We mapped the short reads against the reference genomes using bwa (Li and Durbin, 2009). Mapping results of a representative subset of the samples were checked using *QualiMap* (Garc´ıa-Alcalde *et al.*, 2012).

*Variant analysis.* We used custom analysis scripts to identify mutations that arose and spread during the experiments. Scripts were written in *Python* and use the packages *NumPy*, and *pysam* (Cock *et al.*, 2009; Li *et al.*, 2009; van der Walt *et al.*, 2011).

We used the mapped reads to calculate the number of times each bases ACGT or a gap (-) was observed at every position in every sample (a pile-up). Only positions at which the frequency of a variant changed by at least *>*0.2 and had a coverage of more than a third of the average coverage were considered reliable substitutions. For the preliminary experiment that was sequenced to lower coverage, a minimal frequency change of 0.4 was required.

To find CNVs, we normalized coverage with a position specific average coverage. Coverage of each contig in each run was normalized to the mean coverage along the contig. Then, these normalized coverages were used to calculate the position specific median normalized coverage across all samples. Copy number variations were detected by searching for regions where the normalized coverage dropped below 0.5 fold the position specific median or above 1.8 fold the position specific median. Only regions longer than 200bp were considered.

Regions identified by this criterion were manually inspected for mapping artifacts and temporal signal in the CNV frequency.

## Acknowledgements

We gratefully acknowledge Christa Lanz and Julia Hildebrandt for help with sequencing and Nadine Hoffmann for excellent technical assistance.

## Transparency declarations

All authors declare that no conflicts of interest exist. The funders had no role in study design, data collection and interpretation, or the decision to submit the work for publication.

## Appendix: Preliminary experiment with strain PA77

**FIG. S1.**
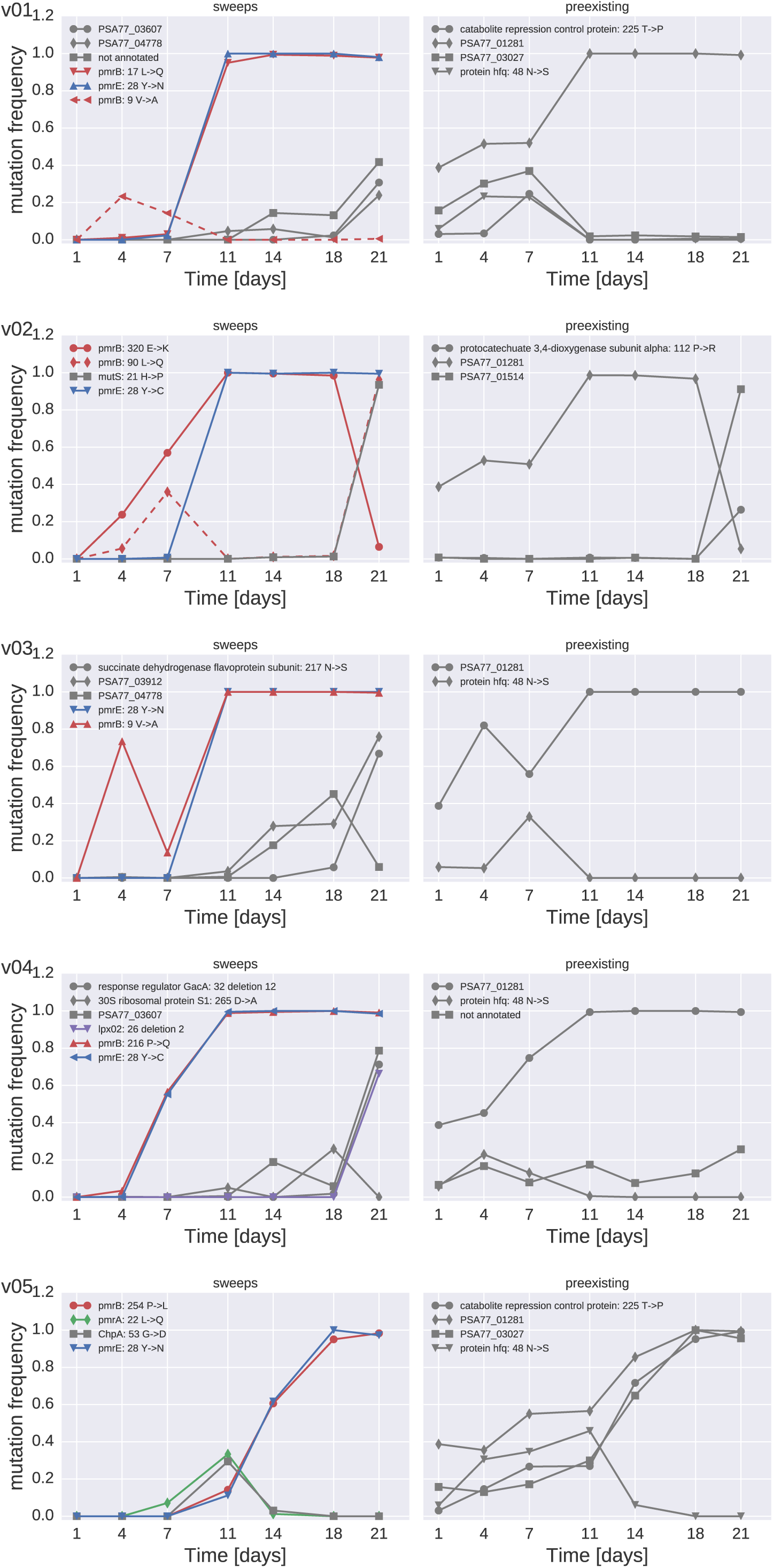
SNP trajectories in cultures of PA77. The left panels contain trajectories of SNPs not observed in the initial sample (sweeps), the right panels contains those already present in the initial sample (preexisting). Mutations that follow the trajectory of the mutator allele are omitted due to their large number. Trajectories of these mutations can be found in Table S7.

**FIG. S2.**
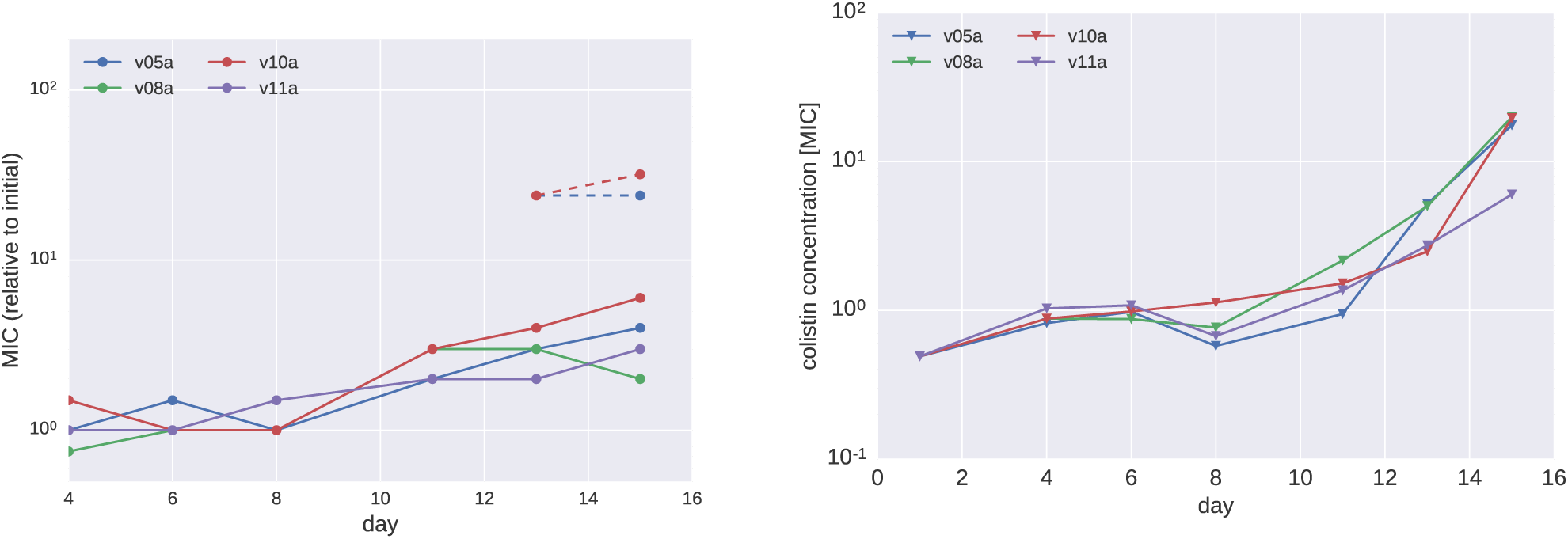
Colistin tolerance in PA77. (A)The time course of MIC in E-tests for each vial in preliminary experiments for strain PA77. Subpopulations showing a higher MIC than the main population were observed after 14 days, respectively. (B) Colistin concentration in each vial in units of the MIC of the initial cultures.

**TABLE S3.** Mutations repeatedly observed in strains PA77. The full list including annotation and locus tag of each mutation in available as supplementary table S3.

| Gene | locus tag | v05a | v08a | v10a | v11a |
| --- | --- | --- | --- | --- | --- |
| PmrB | PSA77_03611 | P169X,M292I | S257N | N41I,P169X | H261Y |
| PmrE | PSA77_06074 | Y28C | Y28C | Y28N | Y28N |
| OstA | PSA77_05098 | Y803X |  |  | L538R |

**FIG. S3.**
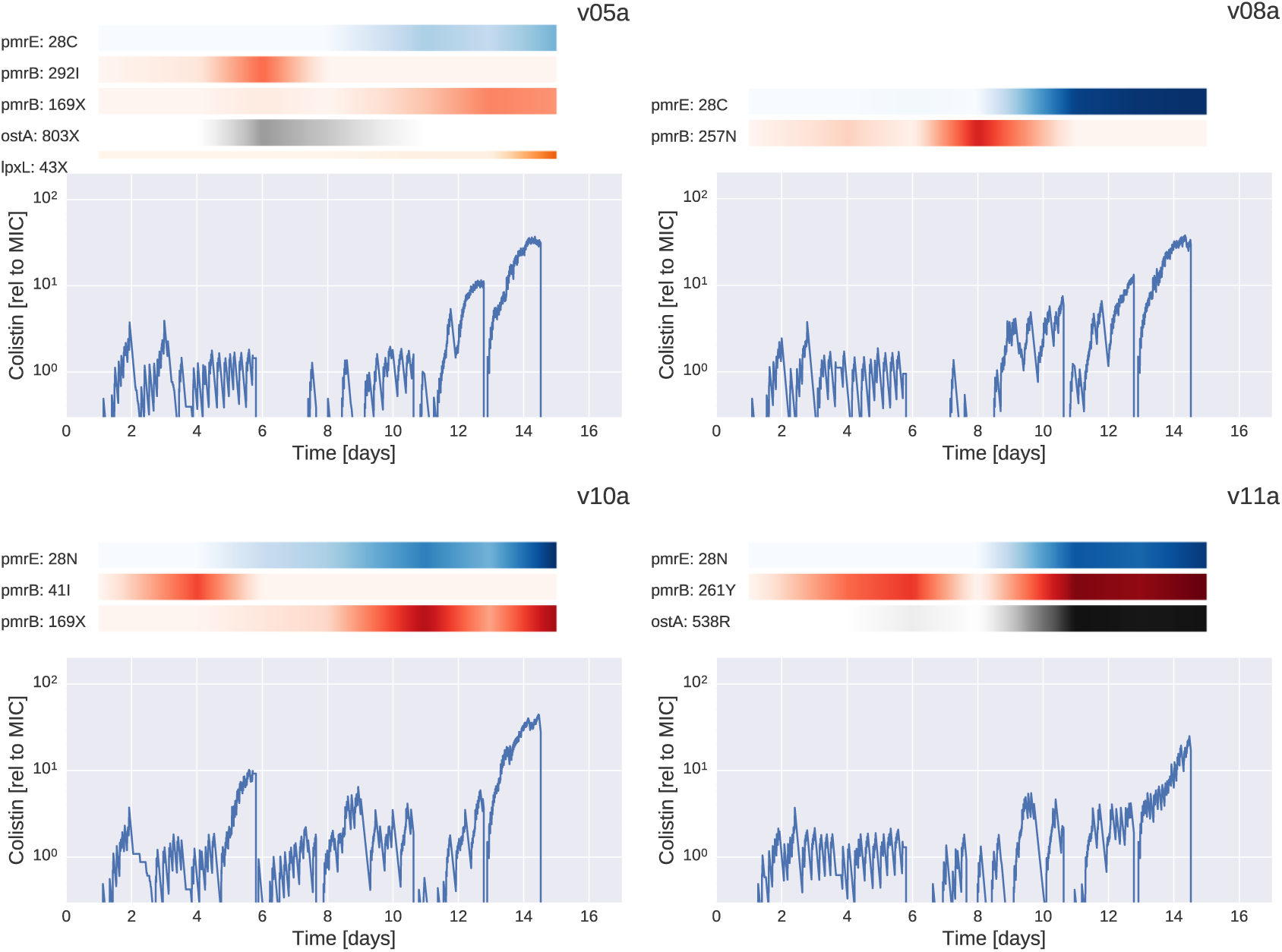
The dynamics of mutations in PA77a. For each culture vial, the plot shows the dynamics of colistin concentration in liquid culture. This concentration is inferred from the cycles of colistin addition and waste removal in 10 minute intervals. The shaded bars above the plots show the abundance of different mutations during the experiment. The frequencies of pmrE and pmrB mutations correlate well with the initial rise in colistin tolerance. The deep dips in colistin concentration every 2-3 days correspond to transfers to fresh culture vials and mark the time points at which samples were taken.

**FIG. S4.**
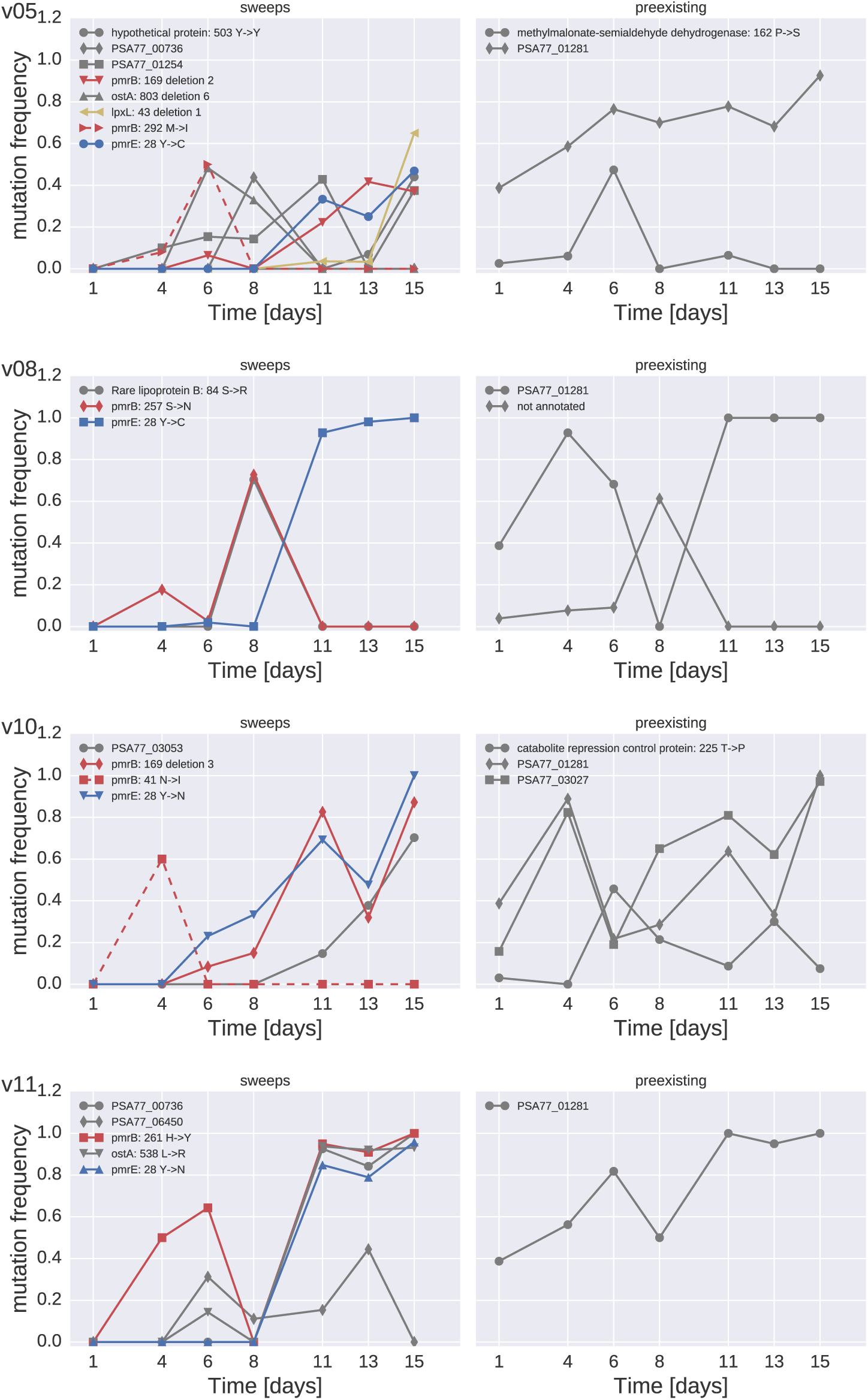
SNP trajectories in cultures of PA77 from the preliminary experiment. The left panels contain trajectories of SNPs not observed in the initial sample (sweeps), the right panels contains those already present in the initial sample (preexisting). This experiment was sequenced at substantially lower coverage than the other two, hence frequency estimates of SNPs are less precise.

## Appendix: Experiment with strain PA83

**FIG. S5.**
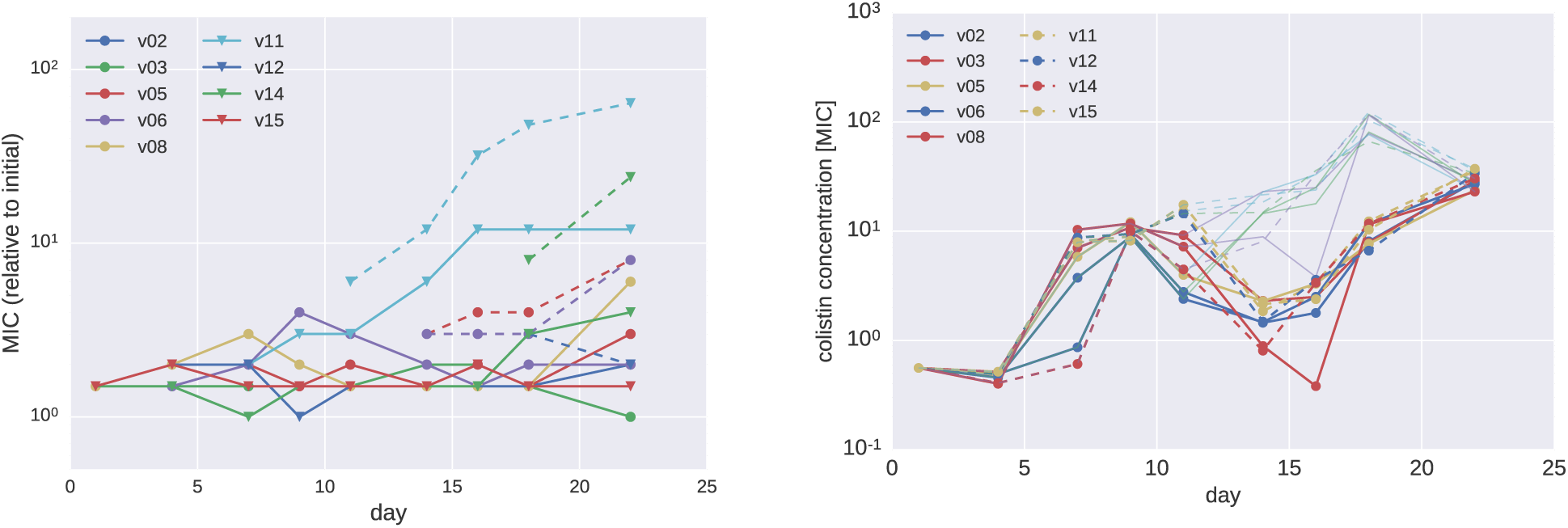
Colistin tolerance in PA83. (A)The time course of MIC in E-tests for each vial in preliminary experiments for strain PA83. Subpopulations showing a higher MIC than the main population were observed after 9 and 14 days, respectively. (B) Colistin concentration in each vial in units of the MIC of the initial cultures. Concentrations for PA83 have at day 14 and 18 have been corrected for an error during the preparation of the colistin stock solution, which had a 10-fold lower concentration than intended.

**TABLE S4.**
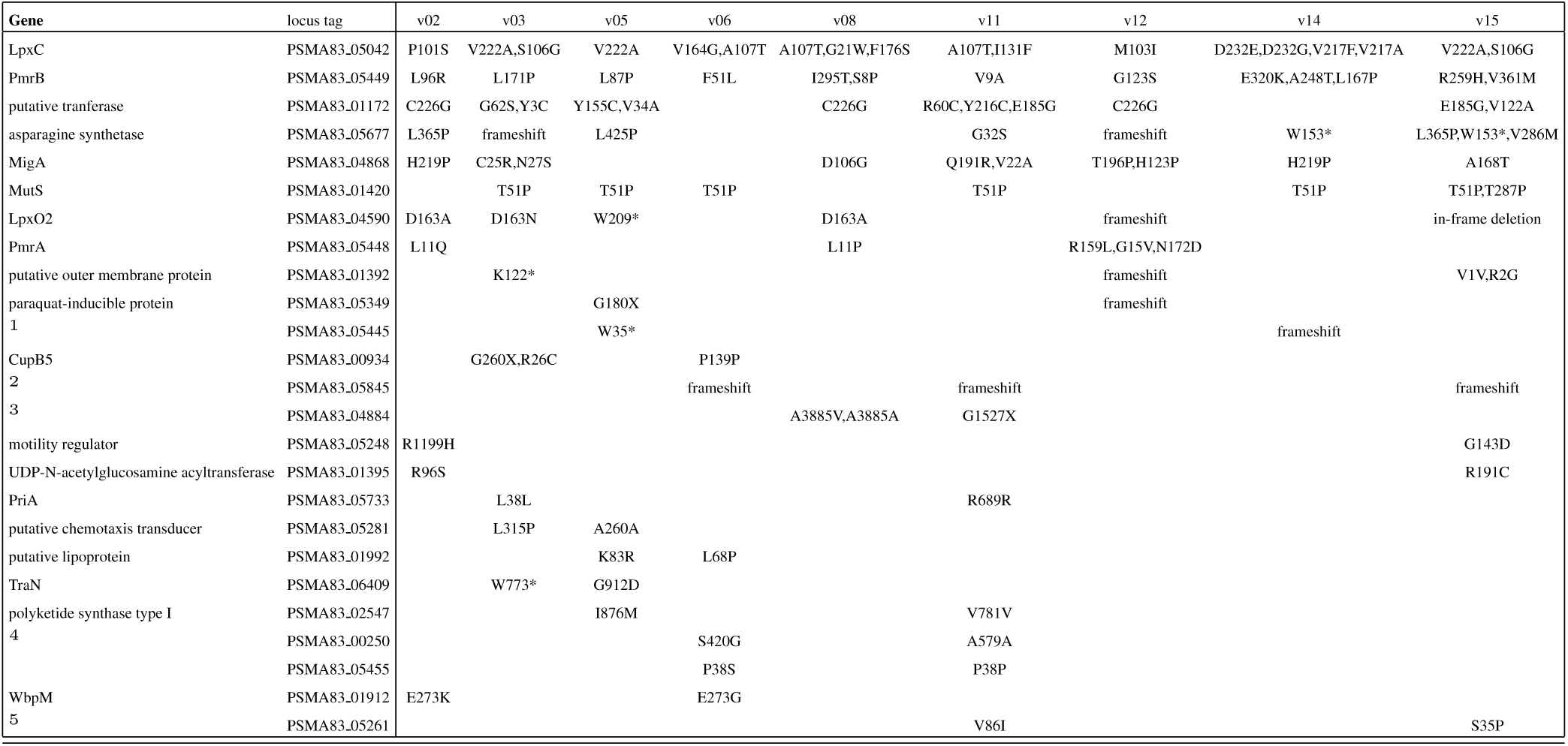
Mutations repeatedly observed in strains PA83. The full list including annotation and locus tag of each mutation in available as supplementary table S4. ^1^ putative S-adenosylmethionine decarboxylase proenzyme; ^2^ putative membrane-bound metallopeptidase; ^3^ filamentous hemagglutinin-like protein; ^4^ putative 4-hydroxyphenylpyruvate dioxygenase; ^5^ large-conductance mechanosensitive channel

**FIG. S6.**
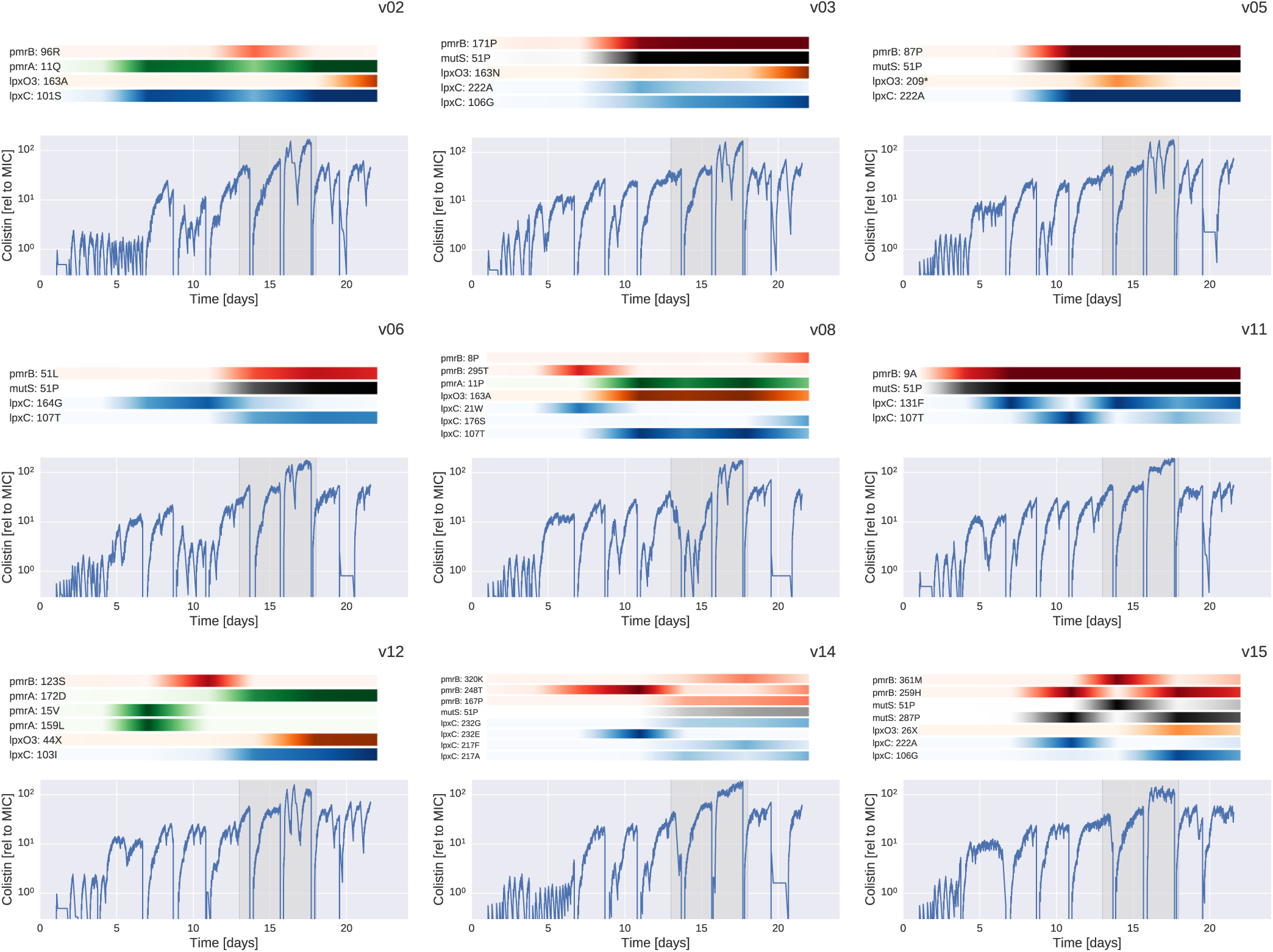
The dynamics of mutations in PA83. For each culture vial, the plot shows the dynamics of colistin concentration in liquid culture. This concentration is inferred from the cycles of colistin addition and waste removal in 10 minute intervals. The shaded bars above the plots show the abundance of different mutations during the experiment. The deep dips in colistin concentration every 2-3 days correspond to transfers to fresh culture vials and mark the time points at which samples were taken. The time interval during which a colistin stock solution with 10-fold lower concentration was used is indicated by a grey box. During these days, the colistin solution in the culture vials was about 10-fold lower than indicated.

**FIG. S7.**
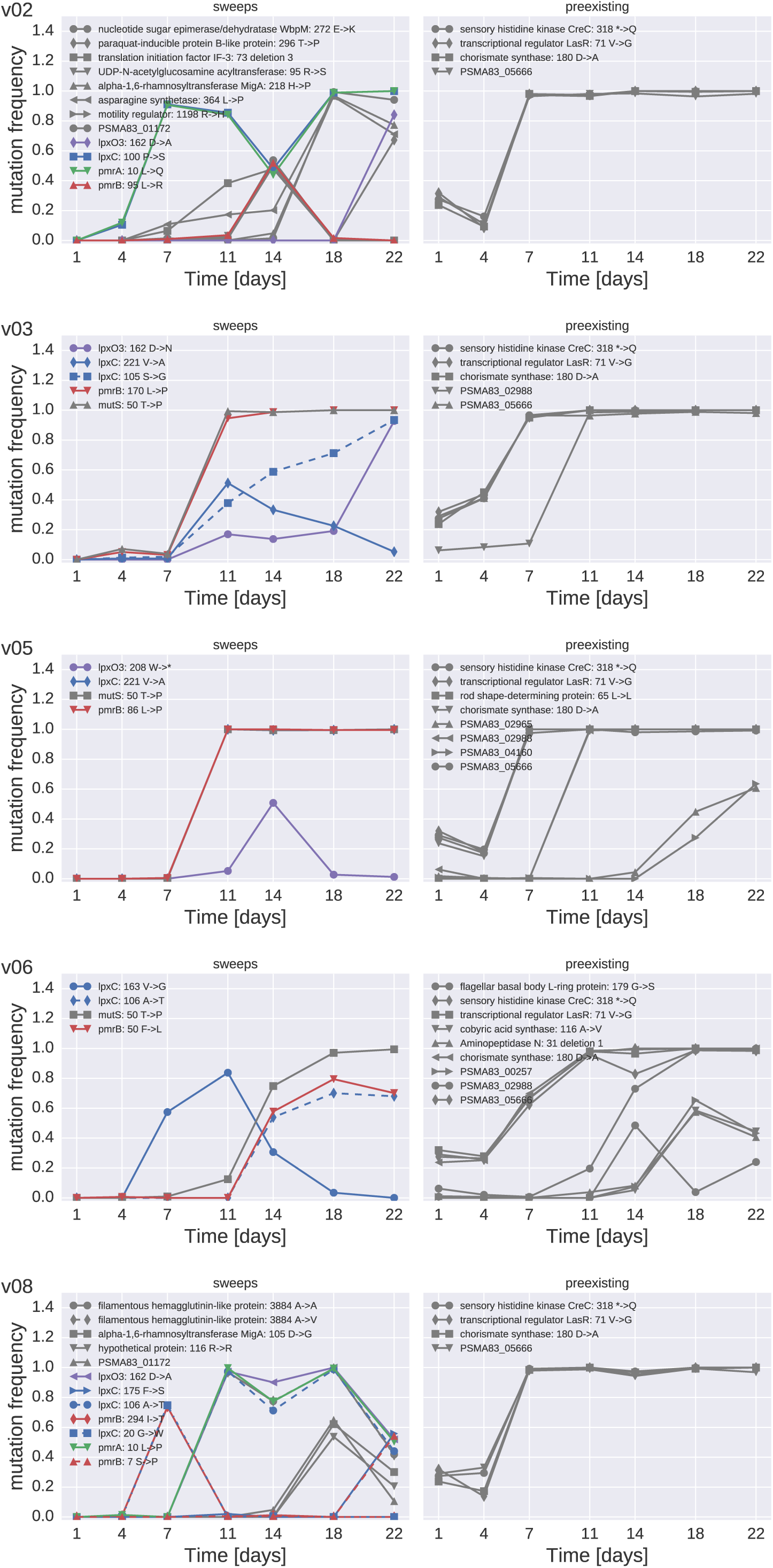
SNP trajectories in cultures of PA83. The left panels contain trajectories of SNPs not observed in the initial sample (sweeps), the right panels contains those already present in the initial sample (preexisting). Mutations that follow the trajectory of the mutator allele are omitted due to their large number. Trajectories of these mutations can be found in Table S7.

**FIG. S8.**
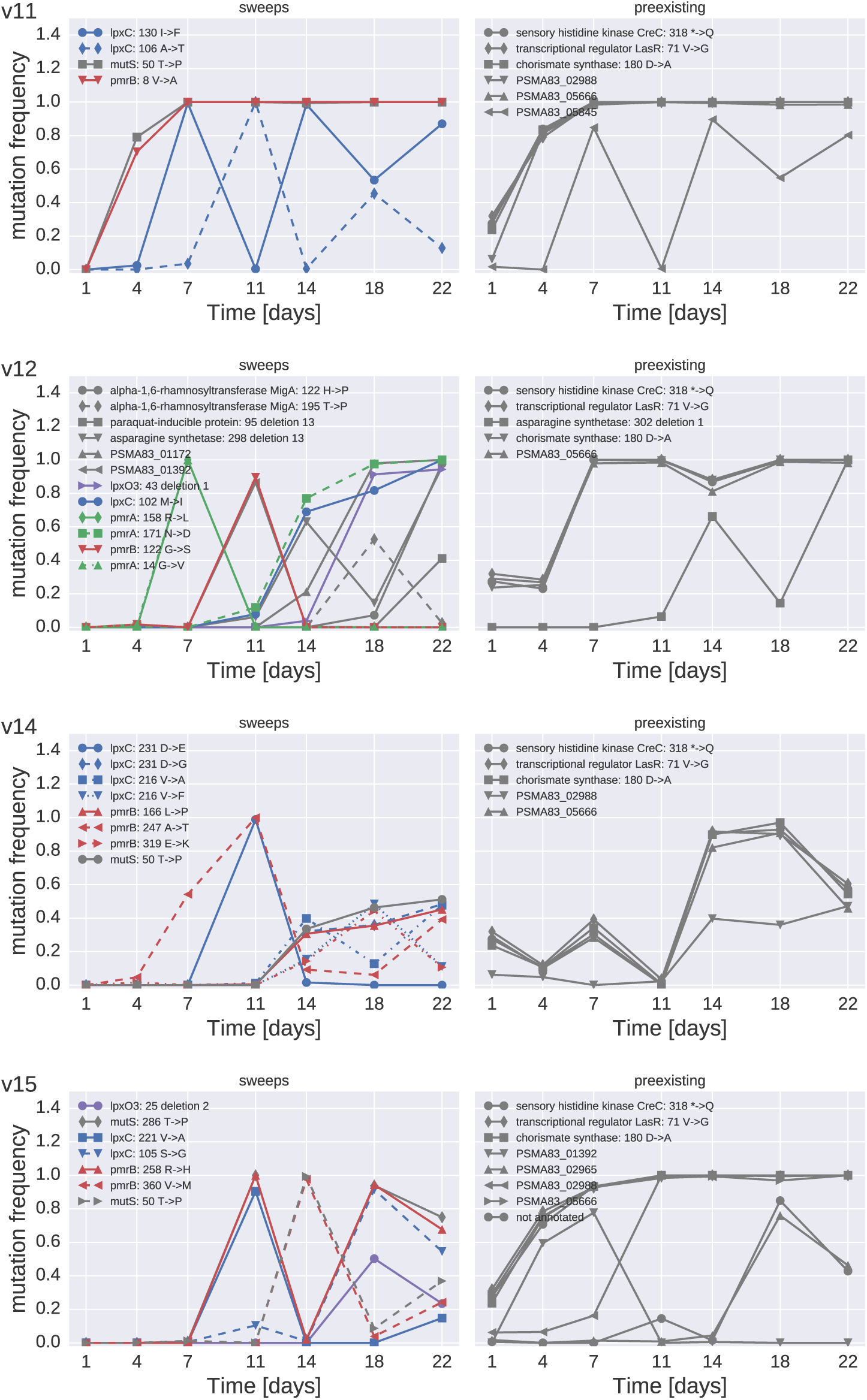
SNP trajectories in cultures of PA83 (contd.). The left panels contain trajectories of SNPs not observed in the initial sample (sweeps), the right panels contains those already present in the initial sample (preexisting). Mutations that follow the trajectory of the mutator allele are omitted due to their large number. Trajectories of these mutations can be found in Table S7.

**FIG. S9.**
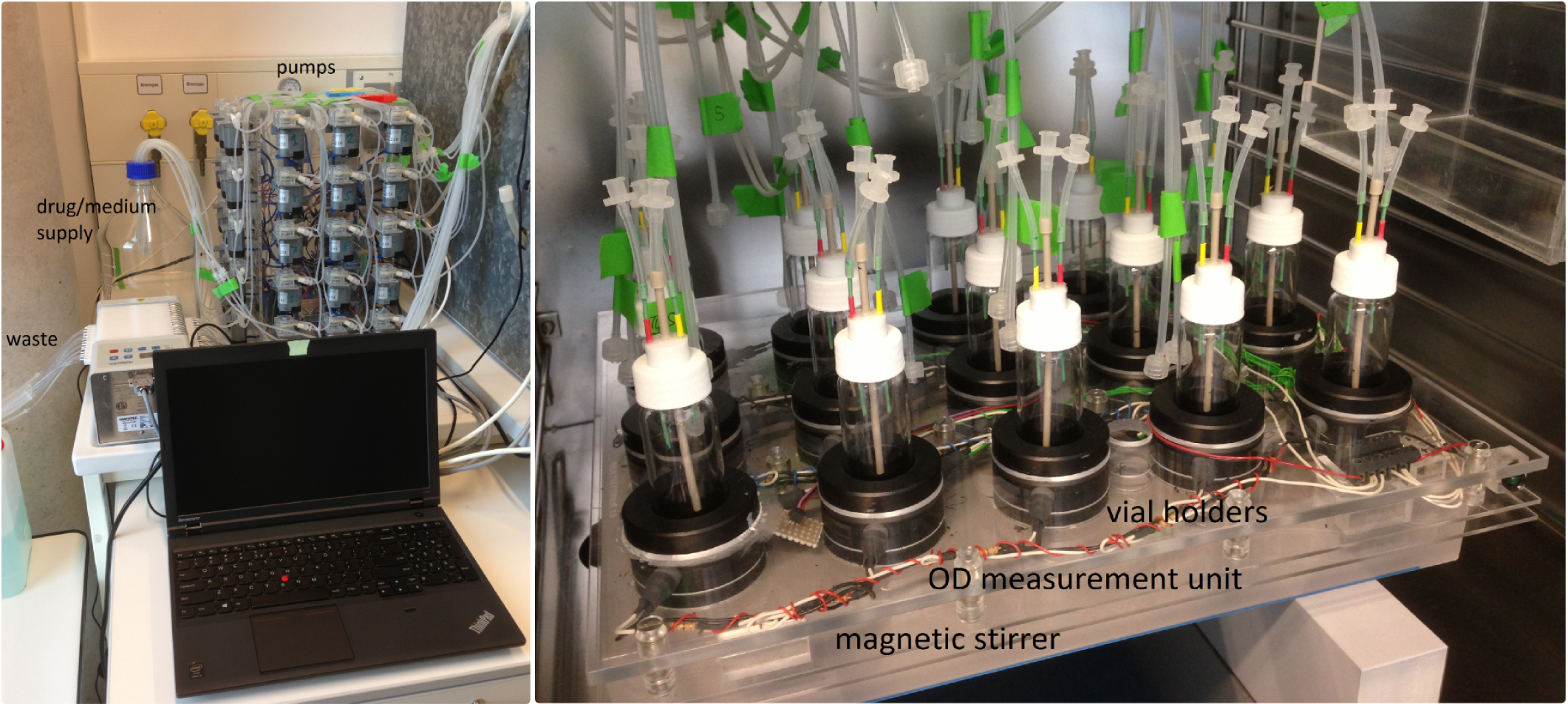
The morbidostat. The left picture shows the pump array, the waste pump, and the inlet in the back of the incubator. The right picture shows the sample holder on a magnetic stirrer and the OD measurement electronics.

## References

Andrews, J. M., 2001, J. Antimicrob. Chemother. 48(suppl 1), 5, ISSN 0305-7453, 1460-2091, URL http://jac.oxfordjournals.org/content/48/suppl_1/5.

Andrews, S., 2015, Fastqc: A quality control tool for high throughput sequence data, http://www.bioinformatics.bbsrc.ac.uk/projects/fastqc/, accessed: Jan 2015.

Aono, R., T. Negishi, K. Aibe, A. Inoue, and K. Horikoshi, 1994, Biosci. Biotechnol. Biochem. 58(7), 1231, ISSN 0916-8451.

Barrick, J. E., D. S. Yu, S. H. Yoon, H. Jeong, T. K. Oh, D. Schneider, R. E. Lenski, and J. F. Kim, 2009, Nature 461(7268), 1243, ISSN 0028-0836, URL http://www.nature.com/nature/journal/v461/n7268/abs/nature08480.html.

Baym, M., T. D. Lieberman, E. D. Kelsic, R. Chait, R. Gross, I. Yelin, and R. Kishony, 2016a, Science 353(6304), 1147.

Baym, M., L. K. Stone, and R. Kishony, 2016b, Science 351(6268), aad3292, ISSN 0036-8075, 1095-9203, URL http://science.sciencemag.org/content/351/6268/aad3292.

Braun, M., and T. J. Silhavy, 2002, Mol. Microbiol. 45(5), 1289, ISSN 0950-382X.

Buchanan, R., N. Stoesser, D. Crook, and I. C. J. W. Bowler, 2014, BMJ Case Rep 2014, ISSN 1757-790X.

Chopra, I., A. J. O’Neill, and K. Miller, 2003, Drug Resist. Updat. 6(3), 137, ISSN 1368-7646.

Cock, P. J. A., T. Antao, J. T. Chang, B. A. Chapman, C. J. Cox, A. Dalke, I. Friedberg, T. Hamelryck, F. Kauff, B. Wilczynski, and M. J. L. d. Hoon, 2009, Bioinformatics 25(11), 1422.

EUCAST, 2016, Recommendations for mic determination of colistin (polymyxin e) as recommended by the joint clsi-eucast polymyxin breakpoints working group, URL http://www.eucast.org/fileadmin/src/media/PDFs/EUCAST_files/General_documents/Recommendations_for_MIC_determination_of_colistin_March_2016.pdf.

Evans, B. A., and S. G. B. Amyes, 2014, Clin. Microbiol. Rev. 27(2), 241, ISSN 0893-8512, 1098-6618, URL http://cmr.asm.org/content/27/2/241.

Fernández, L., C. Álvarez-Ortega, I. Wiegand, J. Olivares, D. Kocíncov, J. S. Lam, J. L. Martínez, and R. E. W. Hancock, 2013, Antimicrob. Agents Chemother. 57(1), 110, ISSN 0066-4804, 1098-6596.

Fernández-Reyes, M., M. Rodríguez-Falcón, C. Chiva, J. Pachón, D. Andreu, and L. Rivas, 2009, Proteomics 9(6), 1632, ISSN 1615-9861.

García-Alcalde, F., K. Okonechnikov, J. Carbonell, L. M. Cruz, S. Götz, S. Tarazona, J. Dopazo, T. F. Meyer, and A. Conesa, 2012, Bioinformatics.

Gould, I. M., and R. Wise, 1985, Lancet 2(8466), 1224, ISSN 0140-6736.

Imamovic, L., and M. O. A. Sommer, 2013, Science Translational Medicine 5(204), 204ra132, ISSN 1946-6234, 1946-6242, URL http://stm.sciencemag.org/content/5/204/204ra132.

Jansen, G., C. Barbosa, and H. Schulenburg, 2013, Drug Resistance Updates 16(6), 96, ISSN 1368-7646, URL http://www.sciencedirect.com/science/article/pii/S1368764614000041.

Jochumsen, N., R. L. Marvig, S. Damkir, R. L. Jensen, W. Paulander, S. Molin, L. Jelsbak, and A. Folkesson, 2016, Nature Communications 7, 13002.

Kang, C.-I., S.-H. Kim, H.-B. Kim, S.-W. Park, Y.-J. Choe, M.-D. Oh, E.-C. Kim, and K.-W. Choe, 2003, Clin. Infect. Dis. 37(6), 745, ISSN 1537-6591.

Katz, D. E., D. Marchaim, M. V. Assous, A. Yinnon, Y. Wiener-Well, and E. Ben-Chetrit, 2016, Int. J. Clin. Pract. ISSN 1742-1241.

Kim, D., S. gun Chung, S. hyup Lee, and J. woo Choi, 2012, African Journal of Microbiology Research 6(21), 4620, ISSN 19960808, URL http://www.academicjournals.org/ajmr/abstracts/abstracts/abstract%202012/9June/Kim%20et%20al.htm.

Krueger, F., 2015, Trimgalore! a wrapper tool around cutadapt and fastqc to consistently apply quality and adapter trimming to fastq files, http://www.bioinformatics.babraham.ac.uk/projects/trimgalore/, accessed: Jan 2015.

Lacour, S., E. Bechet, A. J. Cozzone, I. Mijakovic, and C. Grangeasse, 2008, PLOS ONE 3(8), e3053, ISSN 1932-6203, URL http://journals.plos.org/plosone/article?id=10.1371/journal.pone.0003053.

Leclercq, R., R. Cantón, D. F. J. Brown, C. G. Giske, P. Heisig, A. P. MacGowan, J. W. Mouton, P. Nordmann, A. C. Rodloff, G. M. Rossolini, C. J. Soussy, M. Steinbakk, et al., 2013, Clinical Microbiology and Infection 19(2), 141, ISSN 1198-743X, URL http://www.sciencedirect.com/science/article/pii/S1198743X14602494.

Lee, J.-Y., Y. K. Park, E. S. Chung, I. Y. Na, and K. S. Ko, 2016, Scientific Reports 6, 25543.

Li, H., and R. Durbin, 2009, Bioinformatics 25(14), 1754.

Li, H., B. Handsaker, A. Wysoker, T. Fennell, J. Ruan, N. Homer, G. Marth, G. Abecasis, R. Durbin, and . G. P. D. P. Subgroup, 2009, Bioinformatics 25(16), 2078.

Liu, Y.-Y., Y. Wang, T. R. Walsh, L.-X. Yi, R. Zhang, J. Spencer, Y. Doi, G. Tian, B. Dong, X. Huang, L.-F. Yu, D. Gu, et al., 2016, The Lancet Infectious Diseases 16(2), 161, ISSN 14733099.

López-Rojas, R., J. Domínguez-Herrera, M. J. McConnell, F. Docobo-Perz, Y. Smani, M. Fernández-Reyes, L. Rivas, and J. Pachón, 2011, J. Infect. Dis. 203(4), 545, ISSN 1537-6613.

Magiorakos, A.-P., A. Srinivasan, R. B. Carey, Y. Carmeli, M. E. Falagas, C. G. Giske, S. Harbarth, J. F. Hindler, G. Kahlmeter, B. Olsson-Liljequist, D. L. Paterson, L. B. Rice, et al., 2012, Clin. Microbiol. Infect. 18(3), 268, ISSN 1469-0691.

Malhotra-Kumar, S., B. B. Xavier, A. J. Das, C. Lammens, P. Butaye, and H. Goossens, 2016, Lancet Infect Dis 16(3), 283, ISSN 14744457.

Marvig, R. L., H. K. Johansen, S. Molin, and L. Jelsbak, 2013, PLOS Genet 9(9), e1003741, ISSN 1553-7404.

Moffatt, J. H., M. Harper, P. Harrison, J. D. F. Hale, E. Vinogradov, T. Seemann, R. Henry, B. Crane, F. S. Michael, A. D. Cox, B. Adler, R. L. Nation, et al., 2010, Antimicrob. Agents Chemother. 54(12), 4971, ISSN 0066-4804, 1098-6596.

Moskowitz, S. M., M. K. Brannon, N. Dasgupta, M. Pier, N. Sgambati, A. K. Miller, S. E. Selgrade, S. I. Miller, M. Denton, S. P. Conway, H. K. Johansen, and N. Hiby, 2012, Antimicrob. Agents Chemother. 56(2), 1019, ISSN 0066-4804, 1098-6596.

Moskowitz, S. M., R. K. Ernst, and S. I. Miller, 2004, J. Bacteriol. 186(2), 575, ISSN 0021-9193.

Noteboom, Y., D. S. Y. Ong, E. A. Oostdijk, M. J. Schultz, E. de Jonge, I. Purmer, D. Bergmans, J. W. Fijen, J. Kesecioglu, and M. J. M. Bonten, 2015, Crit. Care Med. 43(12), 2582, ISSN 1530-0293.

Olaitan, A. O., S. Morand, and J.-M. Rolain, 2014, Front Microbiol 5, ISSN 1664-302X.

Pamp, S. J., M. Gjermansen, H. K. Johansen, and T. Tolker-Nielsen, 2008, Mol. Microbiol. 68(1), 223, ISSN 1365-2958.

Pitout, J. D. D., D. B. Gregson, L. Poirel, J.-A. McClure, P. Le, and D. L. Church, 2005, J. Clin. Microbiol. 43(7), 3129, ISSN 0095-1137, 1098-660X, URL http://jcm.asm.org/content/43/7/3129.

Poon, K. K. H., E. L. Westman, E. Vinogradov, S. Jin, and J. S. Lam, 2008, J. Bacteriol. 190(6), 1857, ISSN 1098-5530.

Punina, N. V., N. M. Makridakis, M. A. Remnev, and A. F. Topunov, 2015, Human Genomics 9, 19, ISSN 1479-7364, URL http://dx.doi.org/10.1186/s40246-015-0037-z.

Rodríguez-Martínez, J.-M., L. Poirel, and P. Nordmann, 2009, Antimicrob. Agents Chemother. 53(11), 4783, ISSN 1098-6596.

Sampson, B. A., R. Misra, and S. A. Benson, 1989, Genetics 122(3), 491, ISSN 0016-6731.

Seemann, T., 2014, Bioinformatics 30(14), 2068, ISSN 1367-4811.

Snitkin, E. S., A. M. Zelazny, P. J. Thomas, F. Stock, NISC Comparative Sequencing Program Group, D. K. Henderson, T. N. Palmore, and J. A. Segre, 2012, Sci Transl Med 4(148), 148ra116, ISSN 1946-6242.

Taddei, F., M. Radman, J. Maynard-Smith, B. Toupance, P. H. Gouyon, and B. Godelle, 1997, Nature 387(6634), 700, ISSN 0028-0836.

Toprak, E., A. Veres, J.-B. Michel, R. Chait, D. L. Hartl, and R. Kishony, 2012, Nat Genet 44(1), 101, ISSN 1061-4036, URL http://www.nature.com/ng/journal/v44/n1/full/ng.1034.html.

Toprak, E., A. Veres, S. Yildiz, J. M. Pedraza, R. Chait, J. Paulsson, and R. Kishony, 2013, Nat. Protocols 8(3), 555, ISSN 1754-2189, URL http://www.nature.com/nprot/journal/v8/n3/full/nprot.2013.021.html.

Tumbarello, M., E. Repetto, E. M. Trecarichi, C. Bernardini, G. De Pascale, A. Parisini, M. Rossi, M. P. Molinari, T. Spanu, C. Viscoli, R. Cauda, and M. Bassetti, 2011, Epidemiol. Infect. 139(11), 1740, ISSN 1469-4409.

Tzouvelekis, L. S., A. Markogiannakis, M. Psichogiou, P. T. Tassios, and G. L. Daikos, 2012, Clin. Microbiol. Rev. 25(4), 682, ISSN 0893-8512, 1098-6618, URL http://cmr.asm.org/content/25/4/682.

van der Walt, S., S. Colbert, and G. Varoquaux, 2011, Computing in Science Engineering 13(2), 22.

Willmann, M., A. M. Klimek, W. Vogel, J. Liese, M. Marschal, I. B. Autenrieth, S. Peter, and M. Buhl, 2014, BMC Infectious Diseases 14, 650, ISSN 1471-2334, URL http://dx.doi.org/10.1186/s12879-014-0650-9.

Zanini, F., J. Brodin, L. Thebo, C. Lanz, G. Bratt, J. Albert, and R. A. Neher, 2016, eLife Sciences 4, e11282.

Zankari, E., H. Hasman, S. Cosentino, M. Vestergaard, S. Rasmussen, O. Lund, F. M. Aarestrup, and M. V. Larsen, 2012, J. Antimicrob. Chemother. 67(11), 2640, ISSN 0305-7453, 1460-2091.

